# Transcriptional profiling of early differentiation of primary human mesenchymal stem cells into chondrocytes

**DOI:** 10.1101/2021.08.13.454287

**Authors:** Thomas Schwarzl, Andrea Keogh, Georgina Shaw, Aleksandar Krstic, Elizabeth Clayton, Mary Murphy, Desmond G Higgins, Walter Kolch, Frank Barry

## Abstract

Articular cartilage has only very limited regenerative capacities in humans. Tissue engineering techniques for cartilage damage repair are limited in the production of hyaline cartilage. Mesenchymal stem/stromal cells (MSCs) are multipotent stem cells and can be differentiated into mature cartilage cells, chondrocytes, which could be used for repairing damaged cartilage. Chondrogenesis is a highly complex, relatively inefficient process lasting over 3 weeks *in vitro*. In order to better understand chondrogenic differentiation, especially the commitment phase, we have performed transcriptional profiling of MSC differentiation into chondrocytes from early timepoints starting 15 minutes after induction to 16 hours and fully differentiated chondrocytes at 21 days. Transcriptional gene regulatory networks (GRN) were identified using time-course clustering and upstream-regulator predictions which revealed that cells start epigenetic reprogramming as early as 2 hours after induction and commit to differentiation within 4-6 hours. After that they adapt their gene expression to cater for differentiation specific protein production. These results suggest that interventions to improve the frequency and efficiency of differentiation should target early processes.

## Introduction

Cartilage is a tissue of critical importance in the normal function of the axial skeleton and limb joints, providing the lubricating and shock-absorbing properties of diarthrodial joints and the spine. It also plays a critical role in embryonic limb formation and postnatal growth. Cartilage is a highly specialized avascular and aneural tissue with limited repair capacity (Sherwood et al. 2014). It consists of distinct populations of chondrocytes and an abundant extracellular matrix rich in collagens and proteoglycans (Becerra et al. 2010; Sophia Fox et al. 2009). Traumatic injury of cartilage and degenerative diseases that lead to tissue loss are common and are major causes of disability and loss of mobility. Therapeutic options are limited and there is currently no regenerative treatment that leads to complete restoration of healthy tissue. Thus, there is a compelling need for new therapeutic paradigms and for effective models to understand the regulation of cartilage differentiation and the pathological mechanisms of disease.

Mesenchymal stem/stromal cells (MSCs) represent one such treatment option that has gained much attention in recent years (Zha et al. 2021). In 2018, more than 200 clinical studies using MSCs for cartilage repair were reported (Negoro et al. 2018). Compelling data from preclinical studies and from early stage human trials indicate that MSC treatment is a potential disease-modifying approach (Murphy et al. 2003; Negoro et al. 2018). This often involves the delivery of undifferentiated MSC preparations by intra-articular injection. Autologous chondrocyte (ACI) transplantation is also commonly used as a treatment for acute focal cartilage injury, with good outcomes. This approach has some disadvantages, namely the need to surgically harvest a tissue biopsy from the articular surface for chondrocyte isolation and the limited expansion capacity of primary cultured chondrocytes. MSCs are multipotent progenitor cells isolated from a wide range of adult tissues (Colombini et al. 2019). For therapeutic use the main tissue sources are bone marrow (Prockop 1997), adipose tissue (Prockop 1997; Zuk et al. 2002), umbilical cord (Romanov et al. 2003; Wang et al. 2004) and placenta (Fukuchi et al. 2004). They are readily expandable through many generations in adherent culture using either planar or bioreactor configurations. They possess a well-described trilineage differentiation propensity and can form cartilage, adipose and bone tissues *in vitro* and when transplanted *in vivo* (Dominici et al. 2006). The use of chondrocytes derived from MSCs is an attractive option for cartilage repair treatments and overcomes the obstacles associated with using primary chondrocytes (Pelttari et al. 2008). MSCs are also tolerated as allogeneic therapy, and several advanced therapeutics consisting of allogeneic MSCs have been approved in recent years (Barry 2019; Hoogduijn and Lombardo 2019).

In order to improve the potential therapeutic properties of MSCs for cartilage repair, it is important to understand how and when MSCs commit to chondrogenic differentiation. Of particular interest is the early response to differentiation signals and commitment phase where interventions could be used to increase the number of differentiating cells. Therefore, we examined gene expression during the time-course of chondrogenic differentiation and identified gene regulatory networks (GRNs) that control commitment and differentiation. An obstacle to such studies is the heterogeneity of culture expanded MSC preparations, which consist of mixtures of cells of varying differentiation potential. Therefore, we prepared clonal populations of MSCs from human bone marrow by limiting dilution and obtained a clone with good growth characteristics and clear tri-lineage differentiation potential. Analysis of the GRNs in this clone during chondrogenic differentiation revealed that cells commit to differentiation within 4-6 hours after induction and then adapt their gene expression to cater for differentiation specific protein production.

## Results

A clonal human MSC line was derived from human bone marrow by limiting dilution cloning as described in the Methods section. In total 21 clones were isolated from one donor, of which 11 had trilineage capacity (**Figure 1A**). The selected clone 1F3 retained the full capacity of MSCs for chondrogenic, adipogenic and osteogenic differentiation (**Figure 1B**). Flow cytometric analysis of cell surface antigens showed that these cells were positive for MSC markers CD105, CD90 and CD73 and negative for hematopoietic markers CD45, CD34, CD19, CD3 and CD14, with minor expression of HLA-DR (**Figure 1C**).

**Figure 1.**
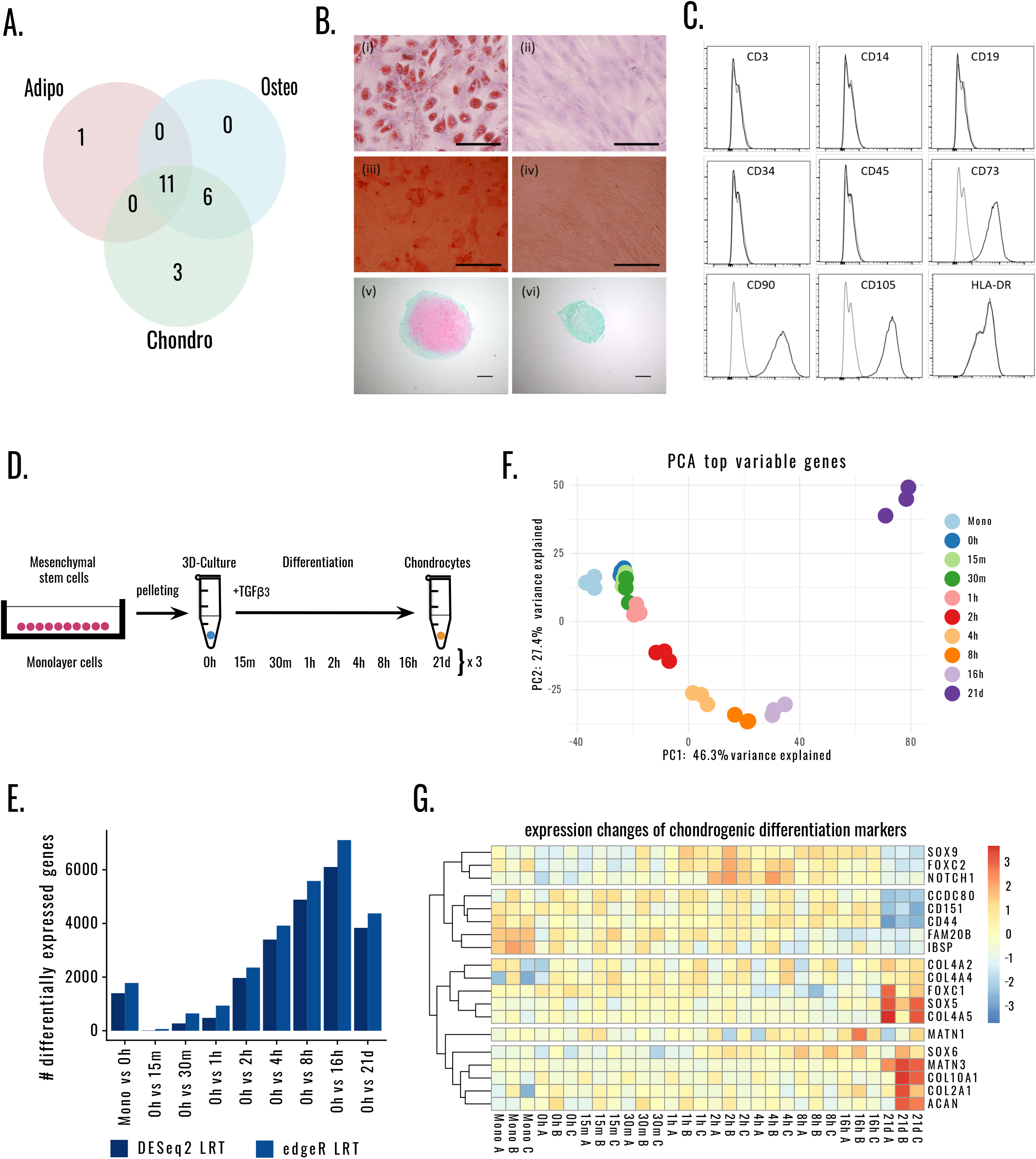
Overview and characterization of a hMSC clone with trilineage differentiation potential. **A**. Schematic of the trilineage differentiation capacity from the bone marrow derived MSC clones from a single donor. **B**. The differentiation assays of the clone 1F3 selected for studying the commitment to chondrogenesis. Adipogenic differentiated (i) and control (ii) wells were stained with Oil Red O, with lipid stained vesicles present in the differentiated wells, scale bar 50μm. Osteogenic differentiated (iii) and control (iv) wells were stained with Alizarin Red, and calcium deposition was noted by positive staining in the differentiated wells, scale bar 200μm. Chondrogenic pellets were stained with Safranin O/Fast Green FCF, with positive staining for glycosaminoglycan seen in the differentiated pellets (v) compared to the control pellets (vi), scale bar 100μM. **C**. Surface profile analysis of clonal cells was carried out using flow cytometry. The clonal population was negative for CD3, CD14, CD19, CD34, CD45 and HLA-DR, but positive for the MSC markers CD73, CD90 and CD105. Isotype controls are shown in grey, and the black line represents each antigen. **D**. Overview of experimental design. Three replicates for Monolayer MSCs, undifferentiated MSCs in 3D culture (0h) and 15m, 30m, 1h, 2h, 4h, 8h, 16h after initiation of differentiation, as well as differentiated chondrocytes (21 days after initiation of differentiation). **E**. Number of differentially expressed genes compared to undifferentiated hMSCs in 3D-culture (pellets, 0h) for DESeq2 and edgeR likelihood ratio tests (LRT). **F**. PCA plot of variance stabilized counts from top 1000 genes. **G**. Differential expression of known markers for chondrogenesis.

To investigate GRNs during chondrogenic differentiation, clone 1F3 cells were induced to differentiate by transferring them from monolayer into 3D culture by gently pelleting the cells and adding TGF-β3. RNA was extracted in triplicates from undifferentiated MSCs in monolayer, undifferentiated MSCs in 3D culture (0h) and 15m, 30m, 1h, 2h, 4h, 8h, 16h after adding TGF-β3, and terminally differentiated chondrocytes (21 days after differentiation) (**Figure 1D**).

Differential gene expression analysis was performed with log likelihood ratio tests in DESeq2 (Love et al. 2014) and edgeR (Robinson et al. 2010), showing that pelleting the cells already had a great impact on gene transcription levels (**Figure 1E**, Mono vs 0h). Principal component analysis (PCA) of the top thousand variable genes suggests that 47.2% of the variance depicted by the first principal component can be attributed to progression in differentiation (**Figure 1F**, x-axis, from monolayer on the left to the 21 d on the right). 27.4% of the variance is associated with early differentiation (y-axis, separation of time-points 0h - 8h). The biological replicates cluster closely together showing that the experimental conditions are very reproducible. An expression table containing all genes is provided in **Supplementary Table S1**. Already 15 minutes after induction of TGF-β3, 11 genes can be detected as differentially expressed (DESeq2, p_adj_-value < 0.05, logFC cutoff +/-0.5, detailed results are provided in **Supplementary Table S2**). The number of differentially expressed (DE) genes increases to 273 genes after 30 minutes, with a rapid further increase of change within the first hours of differentiation. SOX9, NOTCH1, and FOXC2, essential chondrogenic differentiation markers (Gadjanski et al. 2012; Green et al. 2015; Liu et al. 2017), peak at 2 hours after induction by TGF-β3 (**Figure 1G**). In addition to known CD markers, like CD44 and CD151, differential expression of additional CD markers could be observed (**Supplementary Figure 1B**). During differentiation CD99P1, CD109 and CD9 show an increased expression after 8 hours of induction. The expression levels of collagen genes show an alteration after 8 hours and a group of collagen genes including the known differentiation markers COL2A1 and COL10A1 are highly differentially expressed after 21 days (**Figure 1H and Supplementary Figure 1C**) (Charlier et al. 2019). COL4A5 notably correlates with SOX6 expression, whereas COL10A1 shows a correlation with ACAN expression, which is also a marker of mature chondrocytes (Charlier et al. 2019). COL25A1 and COL13A1 are not expressed at 21 days. Human cell markers (Zhang et al. 2019) filtered for stem cell and chondrocyte markers from cartilage and bone marrow tissues, reveal that early differentiation markers like NOTCH1, SOX9, ITGA5, POU5F1 have similar expression profile, as well as NES, NGFR, PECAM1, ITGAV and PDGFRB for later differentiation (**Supplementary Figure 1E**). Chondrocyte markers Aggrecan (ACAN) and Cartilage Oligomeric Matrix Protein (COMP) along with MME, CD4, CD34, COL2A1 are highly differentially expressed in fully differentiated chondrocytes. Stem cell markers like CD44, NT5E and ENG are downregulated after differentiation.

### Functional analysis

Reactome pathway analysis of the DE genes shows the large impact of pelleting on Cell Cycle Checkpoint and signaling pathways, including G-protein-coupled receptor (GPCR) and interleukin signaling (**Figure 2A**). Nuclear Receptor Subfamily 4 Group A Member 3 (NR4A3), a transcriptional activator, and Period Circadian Regulator 1 (PER1), a transcriptional repressor for the circadian clock genes annotated with the enriched Gene Ontology (GO) term “transcription co-factor binding” (GO:0001221, q-value < 0.02) are differentially expressed within the first 15 minutes (**Figure 2B**). Activation of NOTCH signaling and GO terms associated with transcriptional and epigenetic regulation, along with GO terms associated with transcription, can be observed as early as 30 minutes after induction of differentiation (IOD) followed by GPCR signaling from 2 hours (**Figure 2A and B**). Ingenuity Pathway Analysis (IPA) upstream regulator prediction suggests transcriptional changes caused by TGF-β, TNF, PDGF BB, IL1B, and others can be observed as early as 15 minutes after the start of differentiation (**Figure 3A**). Within the first 30 minutes, IPA predicts TGF-β, TNF and other signaling is causing the ‘transactivation of RNA’ along with ‘inhibiting perinatal death and growth failure’ (**Supplementary Figure 2A**). At 1 hour after induction of differentiation, ‘stem cell proliferation’ pathways are activated along with the ‘osteoarthritis pathway’ which expresses SOX9 through TGF-β and maintains ‘differentiation to cartilage’ (**Supplementary Figure 2B and 3**), GO whereas ‘vitamin D3 receptor (VDR/RXR) activation’ is inhibited.

**Figure 2.**
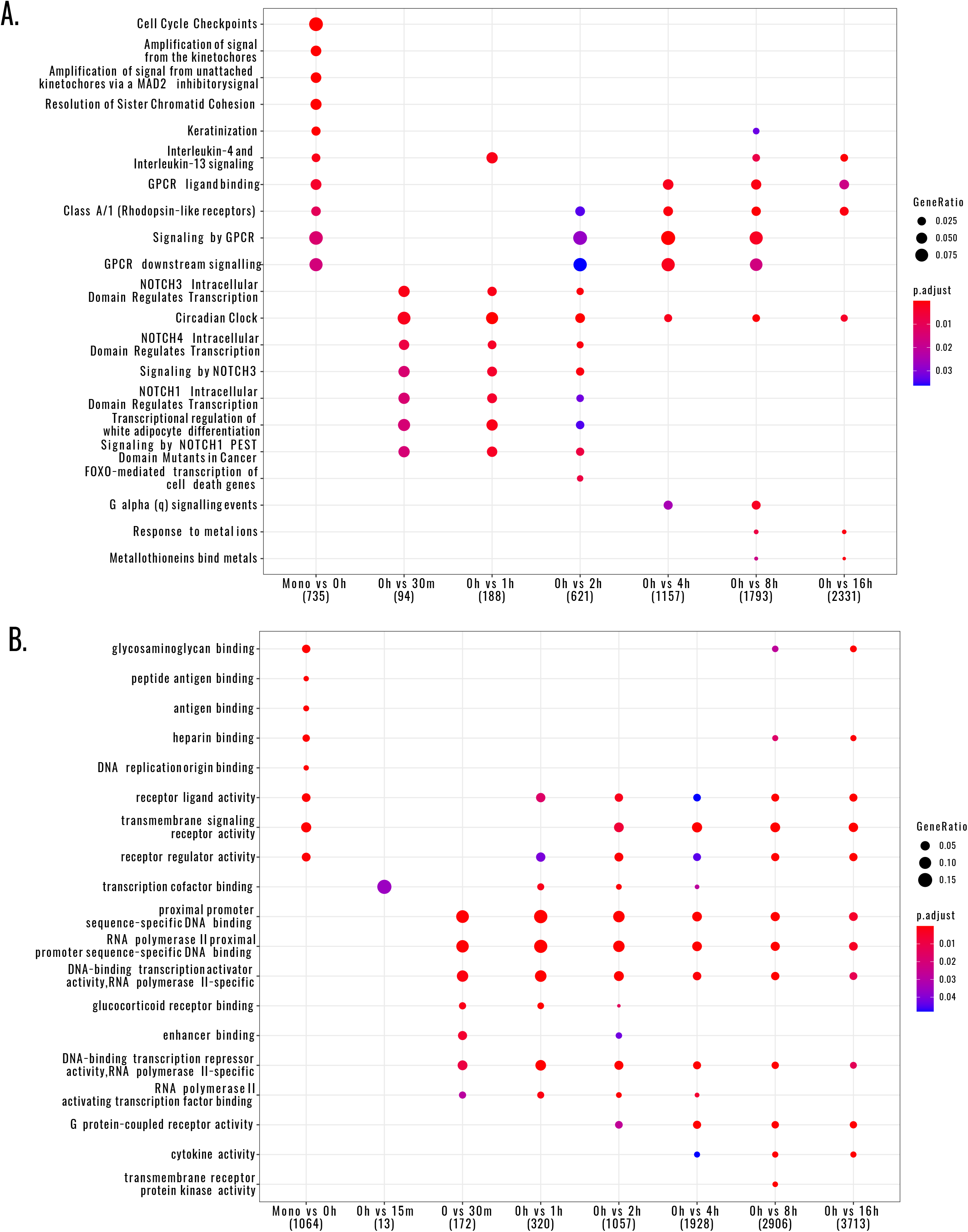
Functional analysis of early differentiation time-points 0h-16h. **A**. Reactome Pathway analysis **B**. Gene Ontology (GO) analysis. The size of the points depicts the gene ratio, how many genes associated with the GO term, or pathway were differentially expressed in the time-point. Color indicated significance, all significant terms or pathways have p_adjusted_ < 0.05.

**Figure 3.**
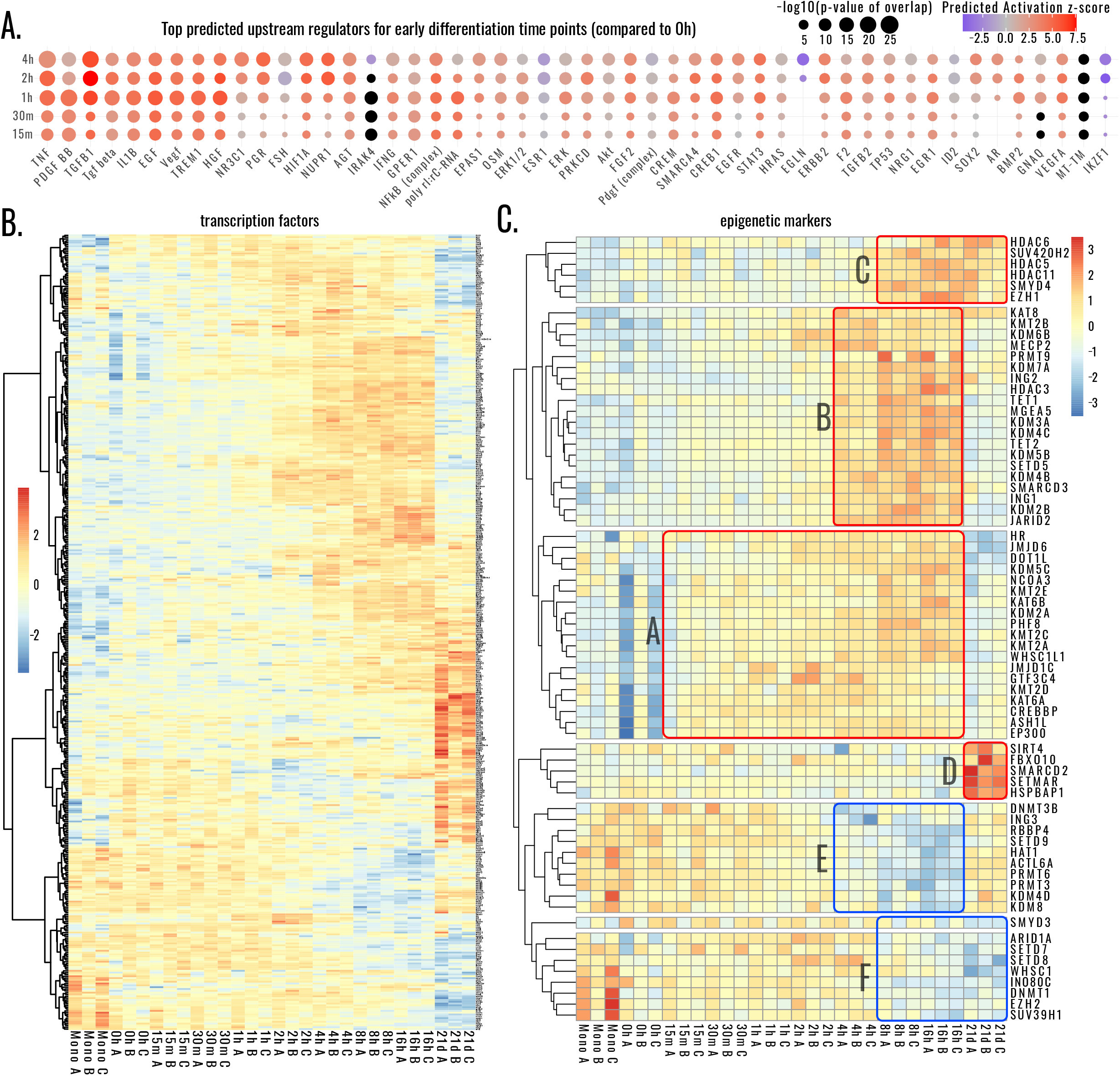
Upstream regulators, Transcription Factors (TF), epigenetic markers. **A**. Top predicted upstream regulators for early differentiation time points by Ingenuity Pathway Analysis. **B**. Heatmaps of differentially expressed TFs and **C**. epigenetic markers (Singh Nanda et al. 2016). Red borders (A-D): upregulated clusters. Blue borders (E-F): downregulated clusters of epigenetic markers.

### Transcription Factors (TFs) and Epigenetic Markers

The expression of the essential pluripotency stem cell TF MYC alongside other TFs is already initiated through pelleting (**Figure 3C**, top, Mono vs 0h). In total 437 TFs are differentially regulated from MSC to differentiated chondrocytes, with waves of TFs expressing over the time-course of differentiation (**Figure 3B**). The expression of epigenetic markers starts as early as 15 minutes after IOD (**Figure 3C**-Cluster A), with clusters peaking at 4 and 8 hours (Cluster B & E and C and F, respectively). Cluster D consisting of SIRT4, FBXO10, HSPBAP1, SETMAR and SMARCD2, is highly expressed in differentiated chondrocytes compared to undifferentiated MSCs.

### Time-course clustering of early differentiation

We clustered differentially expressed genes from early differentiation time-points (0h - 16h) into nine clusters with 1233 genes on average and predicted their upstream regulators using Ingenuity Pathway Analysis (**Figure 4**). Markers for chondrogenesis, MSC chondrogenesis potential, HOX genes and epigenetic regulators were added as annotations for the clusters. Cluster 5 shows upregulation as an immediate response, Cluster 3 peaks at 1-2 hours and then declines, Cluster 4 peaks at 2-4 hours still staying up high at 16 hours, Cluster 2 corresponds to a late response after 4-8 hours, and Cluster 7 represents a very late response after 8-16 hours. Clusters 9, 6, 8 and 1 show a downregulation from early to late timepoints with Clusters 9 and 8 recovering at 16 hours. These clusters contain co-regulated epigenetic regulators and HOX genes, e.g., Cluster 4 includes KDM6B, JMJD6 and other epigenetic regulators which are co-regulated with chondrogenic differentiation markers SOX9, NOTCH1, FOXC2 and CD44. The SOX6 differentiation marker and epigenetic regulators, like JARID2 and Lysine Demethylase (KDM), are co-regulated with the late response of Cluster 2.

**Figure 4:**
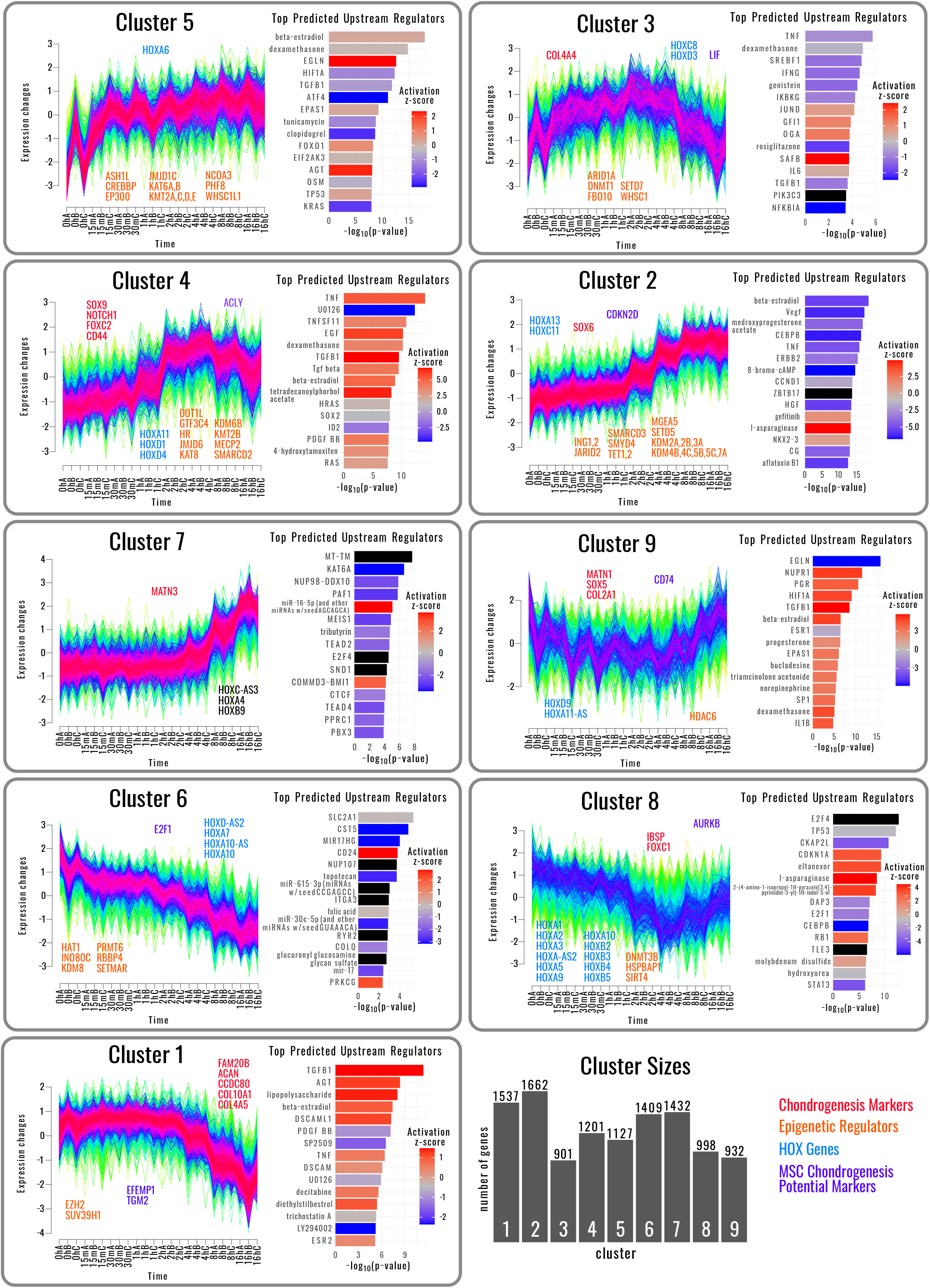
Time-course clusters and upstream regulator prediction of early differentiation time-points 0h-16h and upstream regulator predictions.

Performing upstream regulator and functional analyses on the nine clusters suggested the activation or repression of a combination of transcriptional activators responsible for transcriptional waves during differentiation. The immediate responses of Cluster 5 were predicted to be activated by Egl-9 Family Hypoxia Inducible Factor (EGLN) and a slight deactivation of Hypoxia Inducible Factor 1 Subunit Alpha (HIF1A) as well as the Activating Transcription Factor (AGT), which binds to cAMP response element (CRE). This indicates that the hypoxia pathway becomes activated in the pelleted cells. EGLN is associated with hypoxia and chondrogenic differentiation (Taheem et al. 2018). HIF1A target genes are enriched in early differentiation time-points (**Figure 3**). The micro-RNAs miR-615-3p, miR-30c-5p and miR-16-5p are predicted to contribute to the downregulation of Cluster 6 genes and upregulation of Cluster 7 genes, respectively. The cluster associations are included in **Supplementary Table 1**, and the predicted upstream regulators are provided in **Supplementary Table 3**.

## Discussion

We performed transcriptional profiling for the early differentiation of human MSCs clone 1F3 into chondrocytes using an established differentiation protocol with TGF-β induction of 3D cultured cells (Ng et al. 2008). RNA-seq of monolayer cells, pelleted cells and 15m, 30m, 1h, 2h, 4h, 8h, 16h and differentiated chondrocytes (21days after induction) was used to detect transcriptional patterns and predict GRNs of early differentiation time-points. We were able to detect waves of transactivation of RNAs of epigenetic regulators, TFs, and differentiation markers. Time-course clustering detected 9 clusters of time-course patterns, for which we predicted upstream regulators. The analysis suggests that cell fate reprogramming starts already 15 minutes after induction, and the transcriptional commitment to chondrogenesis is made within the first hours.

Known chondrogenic differentiation markers are a mixture of stem cell and MSC markers which are downregulated in mature chondrocytes (e.g., CD44, CD151, CCDC80, FAM20B) or chondrocyte markers which are upregulated in cartilage cells (e.g., ACAN, COL2A1, COL101, MATN3). Few are indicators of the actual chondrogenic differentiation (like SOX6, SOX9, NOTCH1, FOXC2). In our study, known markers can be observed as expected, which supports, along with experimental validation, the successful chondrogenic differentiation. We provide a thorough functional analysis and annotation of early differentiation for the discovery of novel markers.

Transcriptional activator NR4A3 and repressor PER1 are differentially expressed at 15 minutes after induction. After that, epigenetic markers are differentially expressed in distinct clusters and TFs in waves. We identified previously factors shown to be important for chondrogenesis, such as PDGF, TGF-β, FGF signaling (Massagué and Wotton 2000; Ng et al. 2008). Our analysis also predicted new upstream regulators including TNF-α, HIF1A, IL1B, and others involved in early differentiation. TNF-α stimulation can induce chondrogenesis in ATDC5 cells (Chen et al. 2020), whereas IL1B has been associated with inhibition of differentiation in ATDC5 cells and MSCs (Armbruster et al. 2017; Simsa-Maziel and Monsonego-Ornan 2012). Future studies on the involvement of these regulators in chondrogenesis are therefore recommended.

Out of the early differentiation markers, SOX9 is of great interest, as it was shown to be a target for promoting chondrogenesis (Kim et al. 2018; Ledo et al. 2020; Zhou et al. 2018). Pathway analysis suggests that the TGF-β/Smad signaling pathway leads to an activation of SOX9 expression via the osteoarthritis pathway. Also, the TF KLF15 was shown to increase the chondrogenic differentiation potential of bone marrow MSCs by binding to the SOX9 promoter (Song et al. 2017). In our study, we find KLF15 co-regulated with SOX9, with both peaking at 2 hours after IOD. Furthermore, both TFs co-regulated the earliest chondrogenic differentiation markers NOTCH1 and FOXC2.

Other chondrogenesis relevant genes are HOXC8 and DLX5, which were shown to promote chondrogenesis in human apical papillae derived MSCs (APSCs) by inhibiting LINC01013 (Yang et al. 2020). Our study confirms that HOXC8 and DLX5 increase during chondrogenic differentiation (Cluster 3 and Supplementary Figure 1D). CORIN, a member of the type II transmembrane serine protease class of the trypsin superfamily, was shown to specifically inhibit chondrogenesis in bone marrow MSCs (Zhou et al. 2018). Consistent with these observations, the level of CORIN mRNA is significantly downregulated from 8 hours after IOD.

Vitamin D3 previously was reported to promote chondrogenesis of bone marrow derived MSCs through activation of TGF-β1 signaling (Jiang et al. 2017). On the other hand, it has also been reported to cause chondrocyte hypertrophy and degeneration (Garfinkel et al. 2017). Thus, the role of vitamin D3 is not clear. Our pathway analysis shows an inhibition of vitamin D3 receptors suggesting perhaps that vitamin D3 and TGF-β1 signaling need to be finely coordinated for efficient differentiation to occur.

To date, there are no CD markers known for early chondrogenesis. As expected, the stem cell markers CD44, CD1151 and CCDC80 are downregulated in differentiated chondrocytes, however only few of these CDs are downregulated early in differentiation. These include CD137, a TNF-receptor superfamily member, CD163L1, and CD200 after 8 hours of induction. Further work is required to establish the robustness of such markers.

In summary, our analysis suggests that the commitment of MSCs to chondrogenic differentiation is made remarkably early in the first few hours after induction, and that differentiation itself is executed by coordinated waves of transcriptional activity and epigenetic modifications. Thus, the manipulation of these early events could be a promising way to control the rate and efficiency of chondrogenic differentiation.

## Methods

### Isolation and expansion of bone marrow hMSC clones

hMSCs were isolated from a heparinized bone marrow aspirate harvested from the iliac crest of a healthy 23 year old male volunteer. All procedures were approved by the Clinical Research Ethical Committee of Galway University Hospital and by the National University of Ireland Galway Research Ethics Committee. hMSCs were isolated in vitro by direct plating as described previously(Murphy et al. 2002). Cells were plated at a density of 5×10^4^ mononuclear cells per cm^2^ in alpha MEM (Gibco) supplemented with 10% fetal bovine serum (FBS; Hyclone) and 1ng/mL FGF2 (Peprotech). After five days, when colonies were visible, cells were trypsinized and cloned by limiting dilution into 96 well plates (0.3 cells/well), these clones were termed P1. Wells with single clones were passaged and expanded to P4. The trilineage differentiation potential of the clonal populations were examined, which showed clones with differing propensity for differentiation to the chondrocytic, osteoblastic and adipocytic lineages. A single clone, termed 1F3, with trilineage potential was selected for the study of commitment to chondrogenesis. This clone, as well as having the ability to differentiate to osteoblasts, adipocytes and chondrocytes, also displayed the typical fibroblastic-like morphology and surface marker profile of an MSC.

### Characterisation of MSC clones using trilineage differentiation and surface marker expression

MSC clones were characterised by multi-lineage differentiation and flow cytometry for cell surface marker expression. The multi-lineage differentiation capacity of the MSC clones was assessed by their differentiation to the classical MSC lineages i.e., chondrogenic, osteogenic and adipogenic using established protocols (Murphy et al. 2002).

Chondrogenic potential was examined using a 3-D pellet culture system, where, 2.5×10^5^ cells were pelleted at 100 xg for 5 minutes in complete chondrogenic medium (CCM) consisting of Dulbecco’s Modified Eagle Serum (DMEM, 4.5 g/L glucose), 2 mM glutamine, 100 mM dexamethasone, 50 μg/mL ascorbic acid, 40 μg/mL L-Proline, 1% ITS+ (Insulin, Transferrin, Selenium) (Corning), 1 mM sodium pyruvate and penicillin-streptomycin (100U/mL), supplemented with 10 ng/mL TGF-β3 (Peprotech). Control pellets were incubated without TGF-β3 incomplete chondrogenic medium (ICM). After 21 days, pellets were fixed in 10% neutral buffered formalin, and processed for histology in a Leica ASP300S tissue processor and embedded in paraffin. Sections were stained with Safranin O and Fast Green FCF and imaged with an Olympus BX43 microscope.

For osteogenesis, cells were seeded in culture medium and when the monolayer reached 90% confluence, the medium was replaced with osteogenic medium containing DMEM (1g/L glucose; Sigma Aldrich), 2 mM l-glutamine, 100 nM dexamethasone, 100 μM ascorbic acid, 10 mM β-glycerophosphate, 10% FBS (Hyclone) and penicillin-streptomycin (100 U/ml). Medium was replaced every 3-4 days for up to 14 days, when monolayers were fixed with 10% ice cold methanol, then stained with 2% Alizarin Red and imaged using an Olympus IX71 microscope.

Adipogenesis was assessed by incubating confluent cultures in adipogenic induction medium comprising DMEM (4.5g/L glucose), 2 mM l-glutamine, 10% FBS (Hyclone), 1 μM dexamethasone, 10 μg/mL insulin (Roche), 200 μM indomethacin, 500 μM 3-isolbutyl-1-methylxanthine and penicillin-streptomycin (100U/mL). After 3 days, the culture was transferred to adipogenic maintenance medium comprising DMEM (4.5g/L glucose), 10% FBS (Hyclone), 10 μg/mL insulin (Roche) and penicillin-streptomycin (100 U/mL) for 1 day. This cycle was repeated 3 times after which the cells were maintained in adipogenic maintenance medium for a further 5 days. Cultures were fixed in 10% neutral buffer formalin and stained with Oil Red O before imaging using an inverted Olympus IX71 microscope.

Surface marker expression of MSC clones was carried out by flow cytometry using the BD FACS Canto II flow cytometer (BD Biosciences) using antibodies against CD3, CD14, CD19, CD34, CD45, HLA-DR and the MSC positive markers CD73, CD90 and CD105 (BD Biosciences) as described previously (Ferro et al. 2019). Post-acquisition analysis was carried out using the FlowJo software (Treestar Inc.).

### Study of the commitment towards chondrogenesis

To examine the commitment towards chondrogenesis, clonal cell cultures were switched to a culture medium containing 1% FBS twelve hours prior to chondrogenic induction. Chondrogenic differentiation was carried out in a pellet culture format as described above. Three pellets were set up for each time point. Pellets from the 0h time point remained in ICM, while pellets at 15 min, 30 min, 1, 2, 4, 8 and 16h time points were cultured in CCM. At each time point the 3 pellets were combined into 1 tube, the pooled pellets were washed in PBS and snapped frozen in liquid nitrogen. Clonal cells in monolayer and 21d chondrogenic pellets were included as controls. The time course experiment was carried out three times and each time course served as a technical replica for the experiment.

### RNA extraction and sequencing

Total RNA was extracted using an miRNeasy Mini kit (QIAGEN). The concentration and purity of the isolated RNA was determined using the Nanodrop ND-1000 (Nanodrop Technologies). RNA integrity was measured using the 2100 Bioanalyzer (Agilent Technologies).

Poly(A)+ enrichment to purify mRNA from each sample was performed by hybridization of the RNA with poly(T) oligomers. The mRNA was then converted to cDNA and fragmented, creating cDNA library fragments of approximately 250nt in size. Sequencing adaptors were then added to each cDNA library fragment. The libraries were sequenced on an Illumina GAIIx. The Illumina platform utilizes solid phase amplification on glass flow lanes which have a lawn of primers embedded on them. These primers anneal to the generated cDNA library fragments which were then amplified in clusters. The clusters are then amplified through cyclic reversible termination using fluorescently labelled nucleotides that are imaged as each nucleotide was added. This process generates millions of short (25-300bp) reads, with an average read length of 42bp.

### Bioinformatics Analysis

The raw reads were aligned to Human Gencode (Blanco and Abril 2004) (v23/h38.p3) reference genome with STAR aligner (Dobin et al. 2013) (v2.5.0a) and summarized with featureCounts (Liao et al. 2014) (v.1.6.2). Differentially expressed genes were called with DESeq2 (Love et al. 2014) and edgeR (Robinson et al. 2010), clustering was performed with Mfuzz (Kumar and Futschik 2007) package on variance stabilized, batch effect corrected counts.

The median library size is 21.52 million reads (**Supplementary Figure S1A**). The three differentiated (21-days) samples have 13.69 million reads on average, due to technical challenges of RNA extraction from differentiated chondrocytes.

We have performed log ratio tests on generalized linear models using R/Bioconductor packages edgeR (Robinson et al. 2010) and DESeq2 (Love et al. 2014) with independent hypothesis weighting for multiple hypothesis correction (Ignatiadis et al. 2016). Genes were called significantly differentially expressed if they had absolute log_2_ fold change greater than 1 and p-adjusted-value less than 0.05 in either analysis. Time course clustering was performed for time-points 0h to 16h on variance stabilized, limma-batch corrected counts and then clustered into 9 clusters using the Mfuzz package (Kumar and Futschik 2007).

GO and Pathway analysis were performed with the Bioconductor/R packages ClusterProfiler (Yu et al. 2012) and ReactomePA (Yu et al. 2012; Yu and He 2016). Ingenuity pathway analysis (IPA) was used for upstream regulator analysis and additional pathway analysis (https://www.qiagenbioinformatics.com/products/ingenuitypathway-analysis).

## Supplementary Files

**Supplementary Figure S1. A**. Library sizes of sequencing libraries. **B**. expression changes of differentially regulated CD markers **C**. collagen genes and **D**. DLX. genes **E**. cartilage and bone marrow tissue stem cell and chondrocyte markers.

**Supplementary Figure S2. Ingenuity Pathway Analysis summary. A**. 30 min and **B**. 1h after induction of differentiation

**Supplementary Figure S3. Osteoarthritis Pathway**. Red nodes indicate upregulation in RNA-seq at 1h after induction of differentiation.

**Supplementary Table S1:** Expression Table: This table contains gene annotations, raw counts, library size normalized counts, variance stabilization transformed (DESeq2) counts and variance stabilization transformed counts corrected for gene length. In addition, time-course clustering information and information about differential expression was added.

**Supplementary Table S2:** Table of differentially expressed genes for DESeq2 and edgeR analysis.

**Supplementary Table S3:** Predicted upstream regulators for time-course clusters.

## Declarations

### Ethics approval and consent to participate

All procedures were approved by the Clinical Research Ethical Committee of Galway University Hospital and by the National University of Ireland Galway Research Ethics Committee.

### Consent for publication

Not applicable.

## Availability of data and materials

Raw sequencing data has been submitted to ArrayExpress (E-MTAB-10476), Code and annotations are available in the GIT repository https://git.embl.de/schwarzl/hMSC-diff/, a compiled protocol is provided at http://www.hentze.embl.de/public/MSCdiff/.

## Authors’ Contributions

TS performed the bioinformatic analysis. TS wrote the manuscript. AK and GS contributed sections to the manuscript. WK, DGH and AK edited the manuscript. AK did the IPA analysis of cluster upstream regulators. FB designed the experiment. EC, AnK, GS and MM performed the experimental work. All authors read and approved the final manuscript.

## Acknowledgements

We thank University College Dublin Conway Institute Core Technology Facilities for performing RNA sequencing. We thank Thileepan Sekaran for support with the summarization with featureCounts and Matthias W. Hentze for hosting Supplementary Materials on his group space.

## List of abbreviations

ACAN: Aggrecan
AGT: Activating Transcription Factor
APSCs: human apical papillae derived MSCs
CCM: complete chondrogenic medium
COMP: Cartilage Oligomeric Matrix Protein
CRE: cAMP response element
DE: differentially expressed
DMEM: Dulbecco’s Modified Eagle Serum
EGLN: Egl-9 Family Hypoxia Inducible Factor
FBS: fetal bovine serum
GO: Gene Ontology
HIF1A: Hypoxia Inducible Factor 1 Subunit Alpha
ICM: incomplete chondrogenic medium
IOD: induction of differentiation
ITS+: Insulin, Transferrin, Selenium
KDM: Lysine Demethylase
LRT: likelihood ratio tests
MSCs: Mesenchymal stem/stromal cells
NR4A3: Nuclear Receptor Subfamily 4 Group A Member 3
GPCR: G-protein-coupled receptor
GRNs: gene regulatory networks
PCA: Principal component analysis
TFs: Transcription Factors

