## Supplementary Figure S1-3 for "Transcriptional profiling of early differentiation of primary human mesenchymal stem cells into chondrocytes"

### Supplementary Figure 1

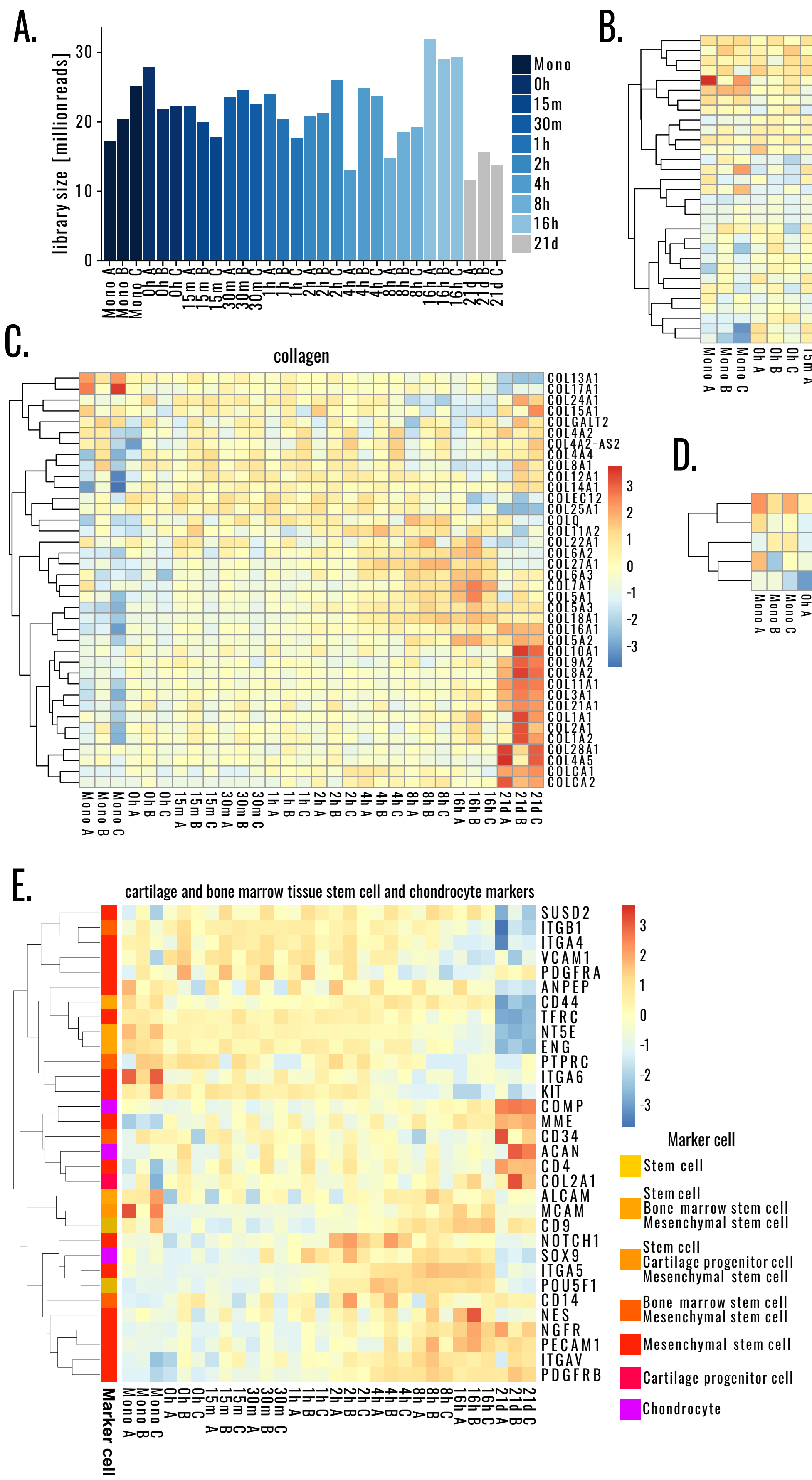

Supplementary Figure 2

A.

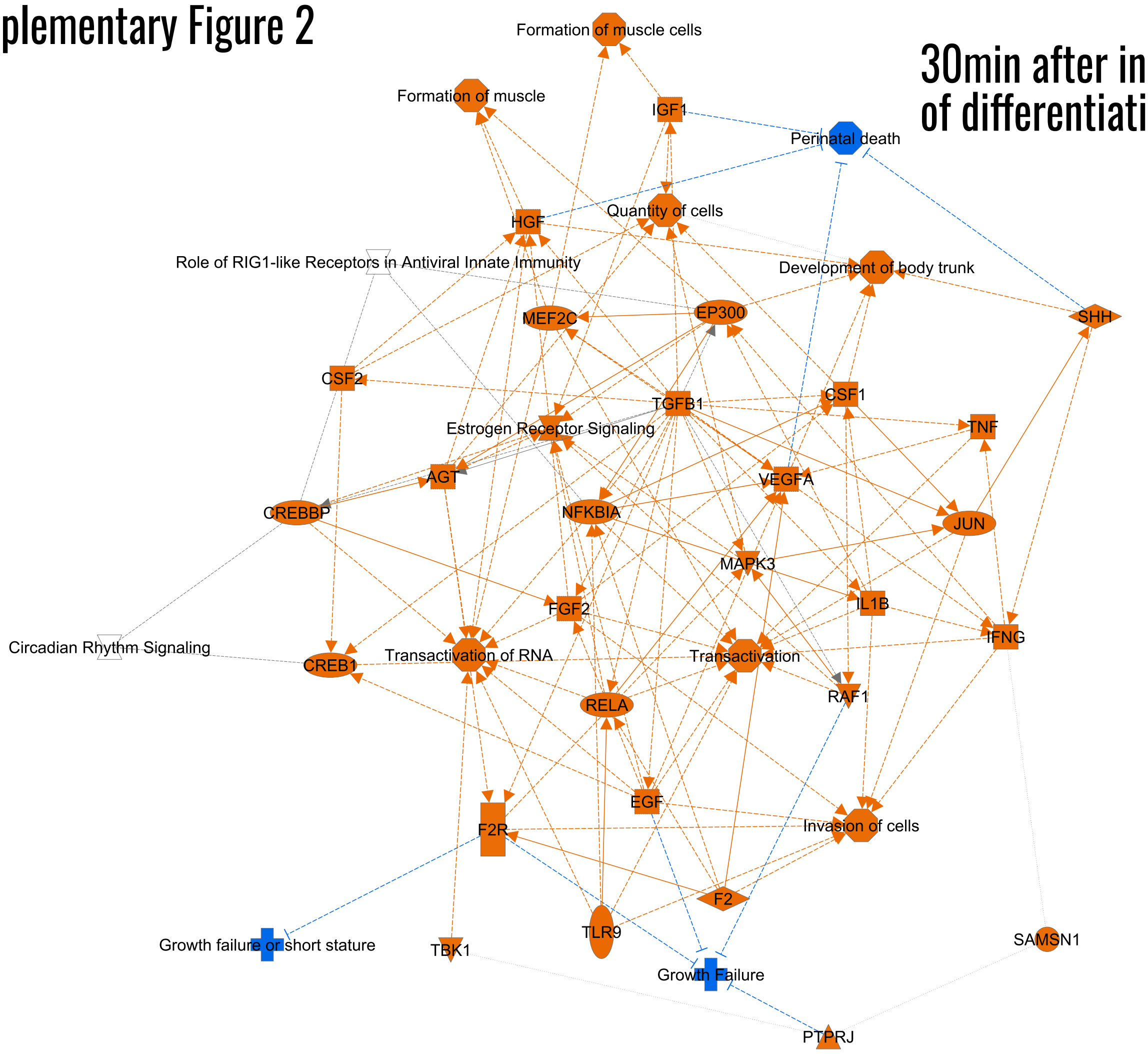


B.

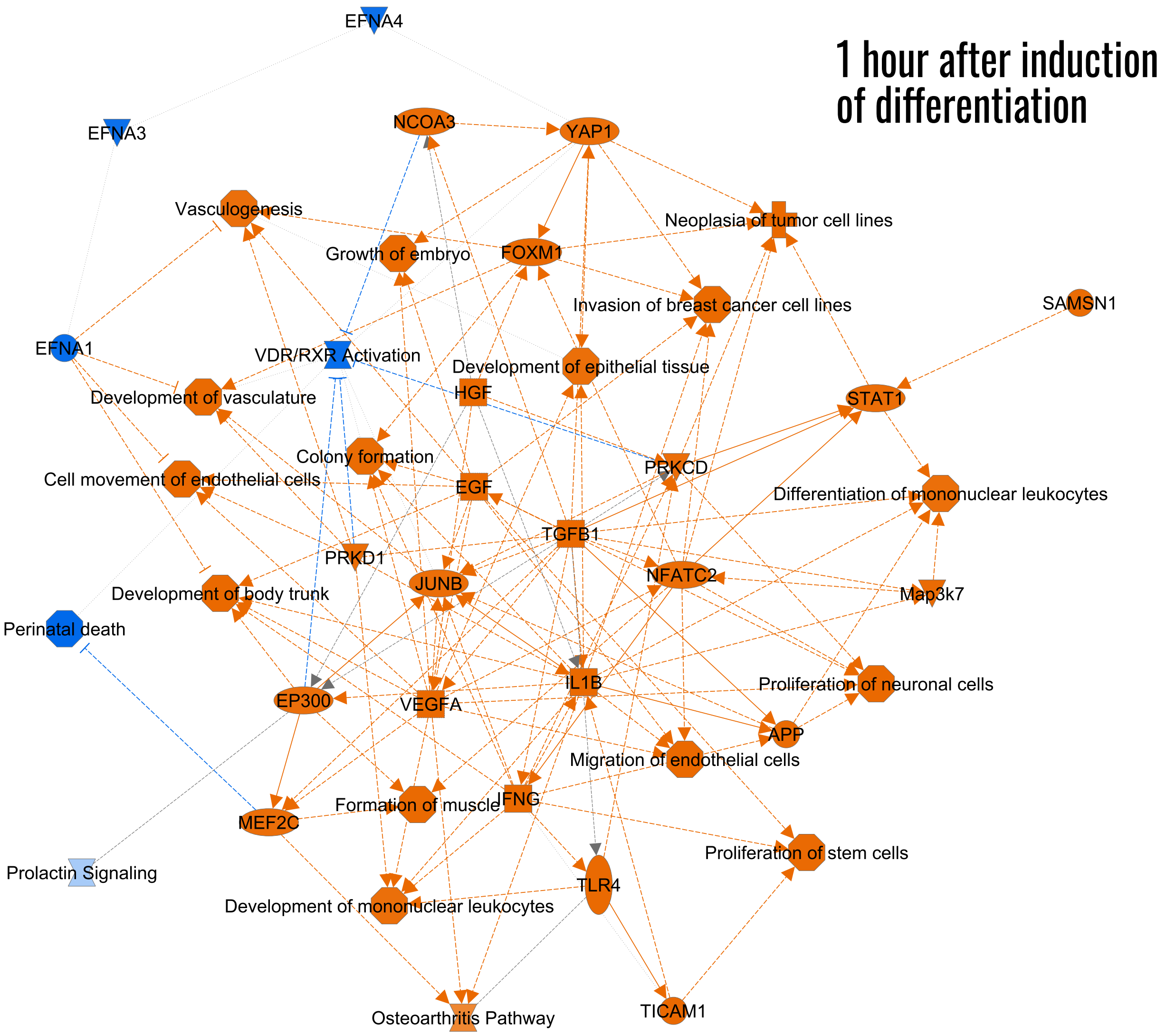


© 2000-2021 QIAGEN. All rights reserved.

### Supplementary Figure 3

#### 1 hour after induction of differentiation

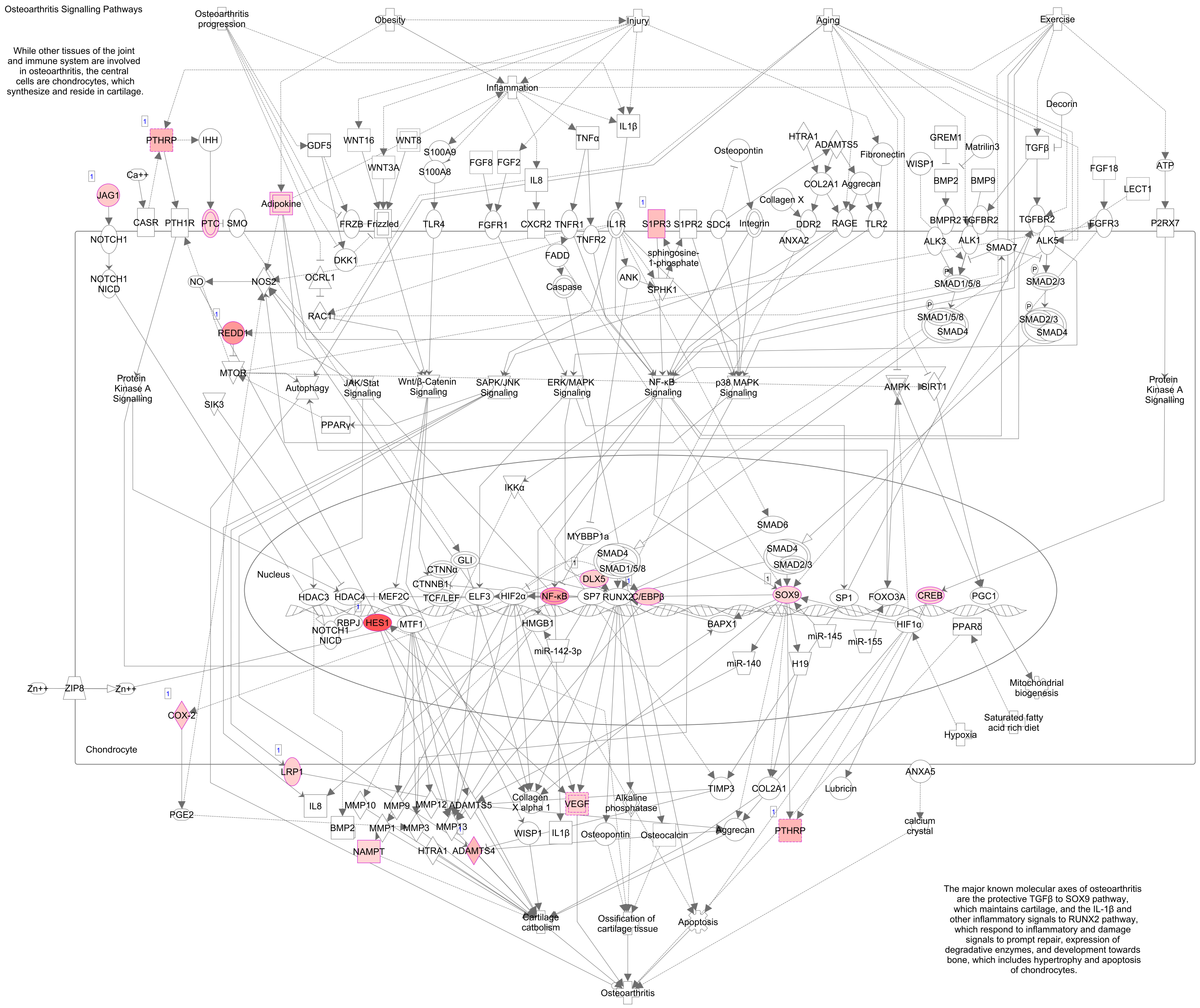


© 2000-2021 QIAGEN. All rights reserved.
